# CUT&RUN detects distinct DNA footprints of RNA polymerase II near the transcription start sites

**DOI:** 10.1101/2020.07.07.191478

**Authors:** Michi Miura, Honglin Chen

## Abstract

CUT&RUN is a powerful tool to study protein-DNA interactions *in vivo*. DNA fragments cleaved by the targeted micrococcal nuclease identify the footprints of DNA-binding proteins on the chromatin. We performed CUT&RUN on human lung carcinoma cell line A549 maintained in a multi-well cell culture plate to profile RNA polymerase II. Long (>270 bp) DNA fragments released by CUT&RUN corresponded to the bimodal peak around the transcription start sites, as previously seen with chromatin immunoprecipitation. However, we found that short (<120 bp) fragments identify a well-defined peak localised at the transcription start sites. This distinct DNA footprint of short fragments, which constituted only about 5% of the total reads, suggests the transient positioning of RNA polymerase II before promoter-proximal pausing, which has not been detected in the physiological settings by standard chromatin immunoprecipitation. We showed that the positioning of the large-size-class DNA footprints around the short-fragment peak was associated with the directionality of transcription, demonstrating the biological significance of distinct CUT&RUN footprints of RNA polymerase II.

## Introduction

CUT&RUN (Cleavage Under Targets and Release Using Nuclease) (Skene and Henikoff 2017) is a powerful tool to map protein-DNA interactions *in vivo*. CUT&RUN directs Protein A/G-fused micrococcal nuclease (pAG-MNase) (Meers et al. 2019a) to the antibody-bound protein-DNA complex, with a subsequent release of DNA fragments cleaved in the presence of calcium ions. There are two major advantages of CUT&RUN over conventional chromatin immunoprecipitation (ChIP). First, CUT&RUN does not require sonication for chromatin fragmentation. Physical shearing of chromatin by sonication does not occur randomly, which can skew signal detection; and excessive sonication may attenuate the epitope of the target protein (Marx 2019). Second, CUT&RUN identifies the footprints of DNA-binding proteins on the chromatin, which revealed distinct binding configurations of transcription factors in yeast (Skene and Henikoff 2017) and mammalian cells (Meers et al. 2019b).

In this study, we performed CUT&RUN on the human lung carcinoma cell line A549 maintained in a multi-well cell culture plate (Fig. 1a) to profile RNA polymerase II (Pol II) around the transcription start sites (TSS). We found that CUT&RUN could validate the bimodal signal around the TSS as previously detected using ChIP (Erickson et al. 2018). However, we also identified a distinct peak at the TSS from short DNA fragments released only by pAG-MNase, suggesting a transient positioning of RNA polymerase II before promoter-proximal pausing, which has eluded detection by conventional ChIP in the steady state of transcription (Erickson et al. 2018).

**Fig. 1.**
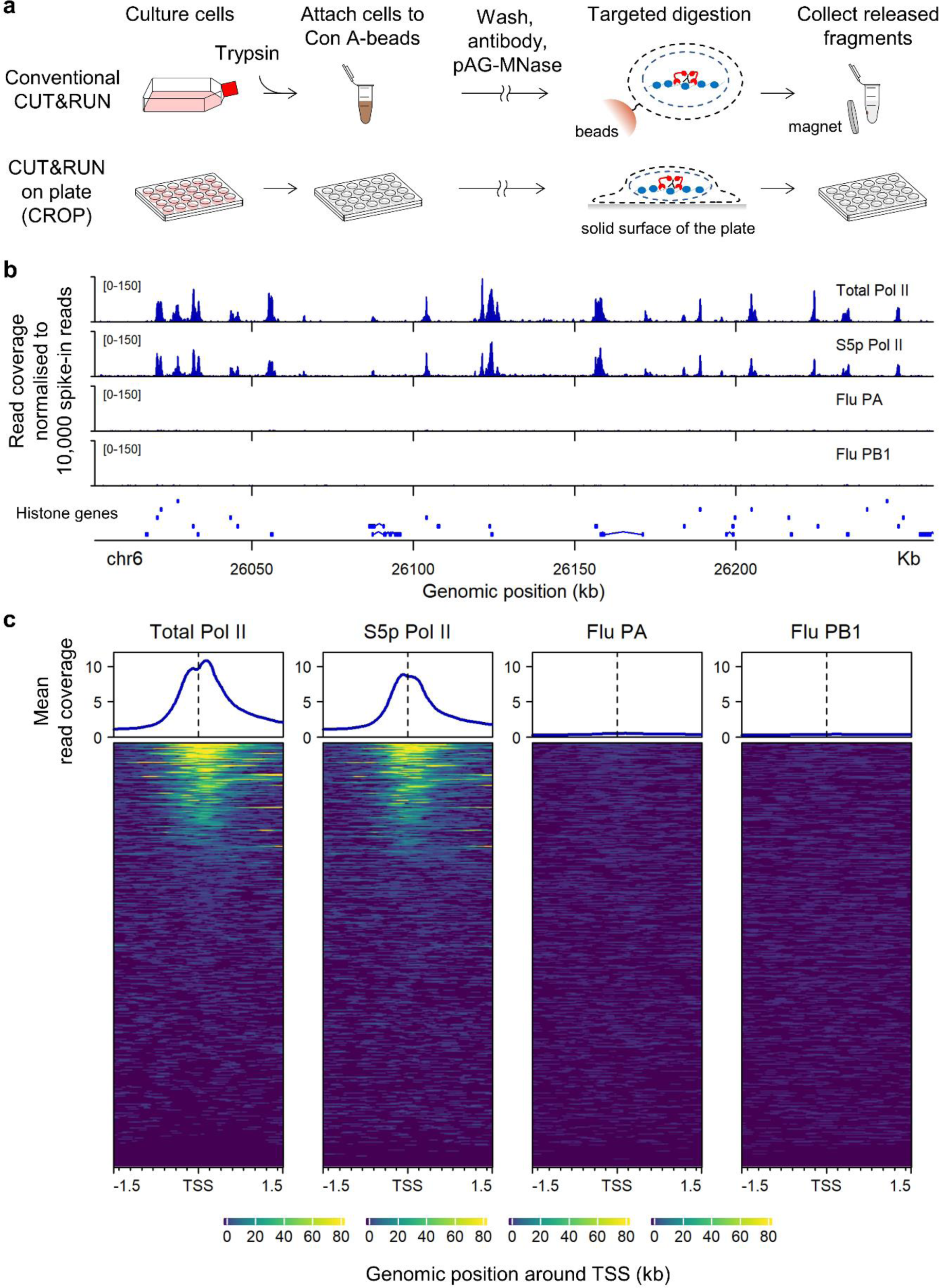
Overall performance of “CUT&RUN on plate”. (a) Protocol overview. In conventional CUT&RUN (top row), cells are harvested and attached on Concanavalin A-magnetic beads. Subsequent cell permeabilisation, antibody binding, and targeted pAG-MNase digestion are performed on these cells immobilised on the beads. The magnetic beads are captured each time the buffers are changed. The alternative approach, “CUT&RUN on plate” (or CROP) (bottom row), circumvents the use of Concanavalin A-magnetic beads. Adherent cells are anchored on the solid surface of the multi-well plate throughout the protocol, enabling rapid buffer change for multiple assays. (b) Pol II signals in the histone gene cluster obtained by CROP. Signals are expressed by the coverage of total aligned reads normalised to 10,000 spike-in read counts. Total Pol II and Serine 5-phosphorylated (S5p) Pol II signals were obtained using the antibodies against RNA polymerase II (Millipore 05-623 and Abcam ab5131, respectively). Flu PA and Flu PB1 are negative controls in which two irrelevant antibodies were used (anti-influenza A PA and anti-influenza A PB1, respectively). (c) Metaplot analysis of the signals from total aligned reads. Spike-in-normalised read coverage was averaged over annotated RefSeq genes that are at least 2 kb away from other genes (n = 13,452). The region within 1.5 kb around the annotated transcription start site (TSS) is presented.

## Results and Discussion

### Overall performance of “CUT&RUN on plate” for profiling Pol II

First, we tested the feasibility of performing CUT&RUN on adherent cells attached to a multi-well polystyrene cell culture plate with Pol II antibodies (Fig. 1a). A549 cells were efficiently permeabilised on the plate with 0.1% Triton X-100 and remained attached throughout the protocol with gentle manipulation. The background cutting was estimated with two irrelevant antibodies (anti-influenza A PA and anti-influenza A PB1). The resulting signals were inspected in the genomic region chr6 p22.2 (Fig. 1b). This location is ideal for efficient visual inspection of Pol II signals because of the compact size and close spacing of multiple histone genes. Our Pol II signals consistently mapped to the transcription units of histone genes with low background as previously reported (Meers et al. 2019a; Skene and Henikoff 2017) (Fig. 1b). The signals within 1.5 kb around the TSS were averaged over “singletons” (genes that are separated from other genes by >2 kb) (Fig. 1c), showing consistent enrichment around the TSS. Therefore, through the successful capture of Pol II signals *in situ* with low background, the performance of our “CUT&RUN on plate” (or CROP) protocol was shown to be comparable with the conventional CUT&RUN (Meers et al. 2019a; Skene and Henikoff 2017).

### CUT&RUN fragments of different length reveal distinct Pol II footprints

The size distribution of aligned paired-end reads showed multiple peaks (Fig. 2a and Online Resource 1). We categorised the aligned reads into four subclasses by length (Fig. 2a), each of which is expected to reflect a unique footprint of the protein-DNA interaction (Meers et al. 2019b; Skene and Henikoff 2017). Indeed, we found that the fragments from different size classes show distinct localisations of Pol II around the TSS (Fig. 2b and Online Resource 2). The signal of the longer reads (>270 bp) conformed to the characteristic bimodal peak as previously reported by ChIP (i.e. the peaks located at either ∼150 bp upstream or downstream of the TSS) (Erickson et al. 2018). By contrast, the shorter fragments (<120 bp) constituted a single narrow peak at the TSS (Fig. 2b). The signals of the shorter (<120 bp) and longer (>270 bp) fragments were mutually exclusive, showing two distinct footprints of Pol II adjacent to each other.

**Fig. 2.**
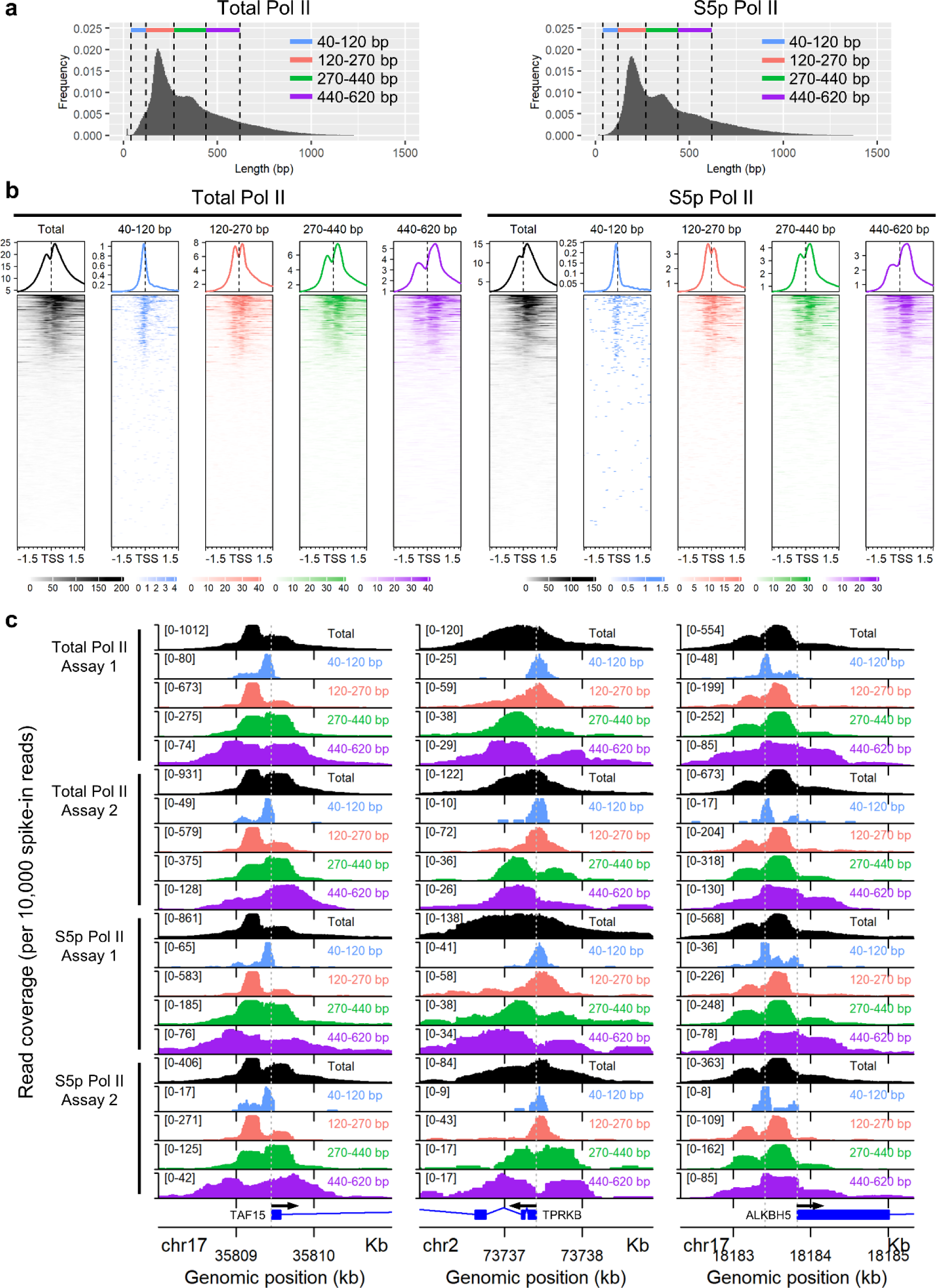
Size-fractionation identifies distinct Pol II footprints. (a) Size distribution of aligned reads (the insert size between the sequence adaptors). The reads were categorised into four subclasses by size, each of which is coded by a unique colour. (b) Total Pol II and S5p Pol II signals from each size class within 1.5 kb around the transcription start site (TSS). The signals in 13,452 annotated RefSeq singleton genes are presented in the heatmap. The metaplot above the heatmap indicates the mean spike-in-normalised read coverage from these 13,452 genes. (c) Total Pol II and S5p Pol II signals from each size classes in individual genes. A 3 kb window of the genomic region around the transcription start site is presented. The vertical axis in each row shows the spike-in-normalised read coverage. The scale is indicated in brackets. The same colour codes are used as per panel (a) to indicate the size classes. The results of four independent assays (two for Total Pol II and the other two S5p Pol II) are shown for each gene. The grey dashed line indicates the annotated transcription start site. An additional line is shown in ALKBH5 to indicate the deviation of the short-fragment peak from the TSS.

Recent studies using ChIP-seq, as well as nascent transcript-sequencing, have shown that Pol II is paused at ∼50 bp downstream of the TSS (Core et al. 2008; Erickson et al. 2018; Kwak et al. 2013; Nojima et al. 2015; Shao and Zeitlinger 2017). Since the duration of Pol II in the promoter-proximal pausing state is about ten times longer than that of Pol II in the pre-initiation state (Darzacq et al. 2007; Steurer et al. 2018), a majority of reads obtained by ChIP is from the paused site, approximately 50 bp downstream of the TSS. Because ChIP fragments are typically hundreds of base pairs long, the rare signal of the poised Pol II at the TSS is often obscured. While Erickson *et al*. had previously identified the signal of the poised Pol II at the TSS by ChIP, this signal was only exposed following the depletion of Pol II in hypotonic condition with the use of triptolide to block the transcription initiation (Erickson et al. 2018). In contrast, we accurately identified the unique positioning of Pol II at the TSS in the physiological condition through the recovery of its unique footprint.

### Skewed positioning of the large-size-class Pol II footprints is associated with the directionality of transcription

As expected, the bimodal signal of the longer fragments (>270 bp) is skewed towards the region downstream of the TSS (Fig. 2b) (Erickson et al. 2018); when genes were inspected individually, however, we observed a variation in the skewness of large-size-class Pol II footprints around the TSS (Fig. 2c). This observation prompted us to stratify genes by the pattern of the large-size-class Pol II footprints.

We observed slight deviation of the short-fragment peak from the annotated transcription start site in some genes (e.g. ALKBH5 in Fig. 2c), possibly due to the alternative transcription start site. Therefore, we quantified the read coverage of the large-size-class fragments in 500 bp upstream or downstream from the short-fragment peak for each gene (Fig. 3a). We selected genes with the short-fragment peak above the background and identified the peak position by local polynomial fitting (Fig. 3b). Genes which had the peak within 500 bp from the annotated transcription start site were included in this analysis (Fig. 3b). The signals were quantified for each of the two size classes (120-270 bp and 270-440 bp), and the skewness of the footprint is expressed by the ratio of the downstream signal to the upstream signal (Fig. 3a). The two size classes showed similar skewness for each gene (Fig. 3c and Online Resource 3). About 14% of the genes were positively skewed (skewness ≥ 1 for both two size classes) and 8% negatively skewed (skewness ≤ −1 for both two size classes) (Fig. 3c and Online Resource 3).

**Fig. 3.**
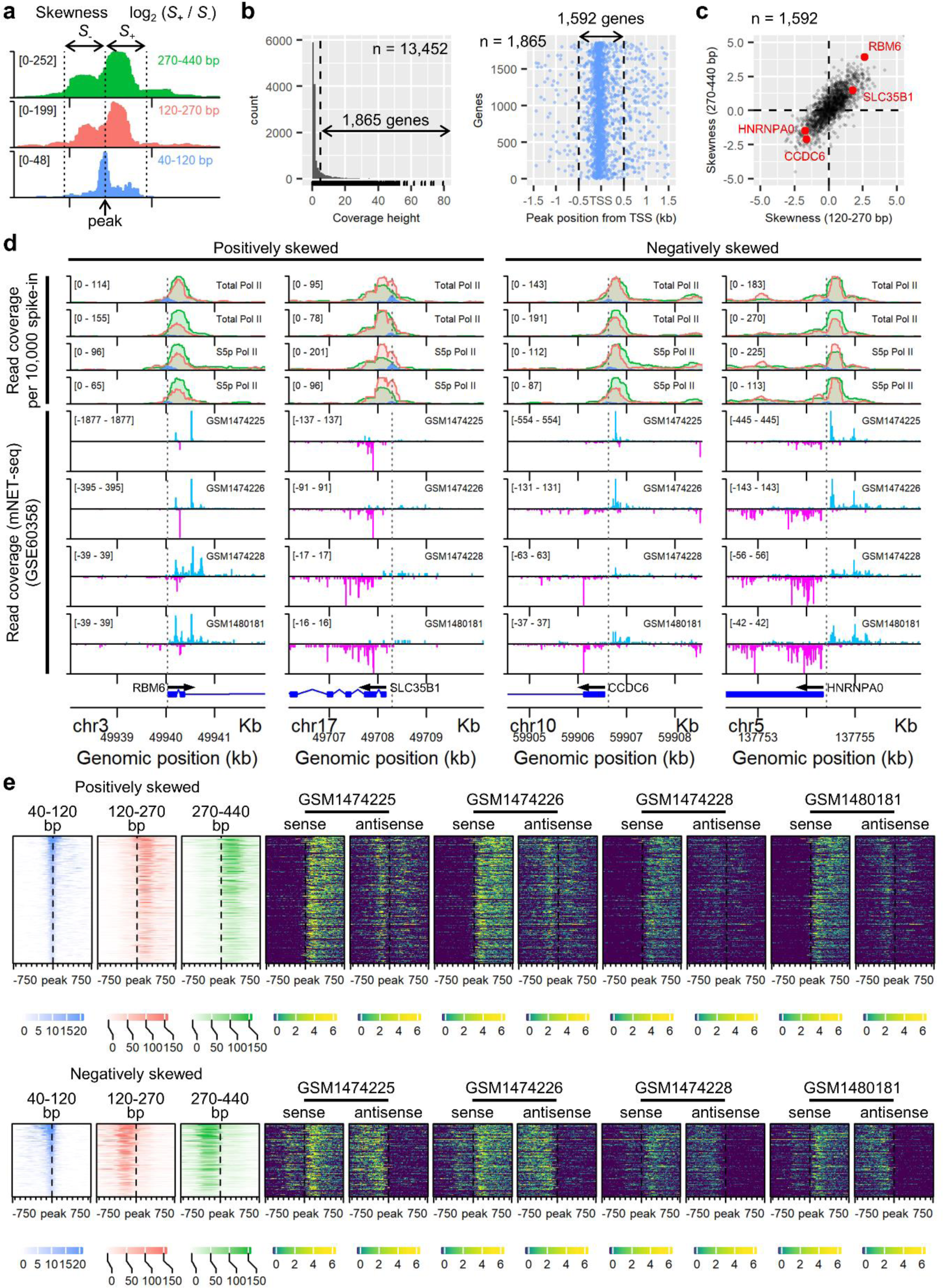
Skewed distribution of the large-sized Pol II footprints around the TSS is associated with the directionality of transcription. (a) Schematic diagram of quantifying the skewness of the Pol II signals at the TSS. The position of the short-fragment peak was identified and then the painted area (i.e. the read coverage) in 500 bp upstream from the peak (denoted as S-) and 500 bp downstream from the peak (S+) was calculated. Skewness is expressed by the logarithmic ratio of S+ to S-in this study. (b) (left) Distribution of the hight of the short-fragment coverage from the 13,452 singleton genes. Genes that had the short-fragment signal above the background (dashed line) were selected to allow robust identification of the short-fragment peak (1,865 genes). (right) The position of the short-fragment peak relative to the annotated transcription start site for each gene. The vertical axis indicates the index of 1,865 genes. Genes that had the peak position within 500 bp around the annotated transcription start site were further selected (1,592 genes). (c) Distribution of the skewness from 1,592 genes. The skewness for the two size classes is plotted for each gene (120-270 bp on the x-axis and 270-440 bp on the y-axis). Four genes that were recurrently identified as skewed, either positively or negatively, are indicated in red. (d) DNA footprints of RNA polymerase II identified by CUT&RUN in this study, and the nascent transcript-sequencing data from the public database. The alignment is shown near the TSS of the four genes from panel (c). The first four rows in each column show CUT&RUN signals from three size classes (40-120 bp, 120-270 bp and 270-440 bp). The same colour code is used as per Fig. 2a to indicate the size classes. The signals were normalised to 10,000 spike-in reads. The scale of the vertical axis in each panel is indicated in bracket. The second four rows show nascent RNA transcription from Gene Expression Omnibus (GSE60358), which contains four mNET-seq assays. The signal of the plus-strand-nascent transcripts is indicated in light blue, and the minus-strand-transcripts in magenta, regardless of the gene orientation shown at the bottom of each column. The grey dashed line indicates the position of the short-fragment peak. (e) Heatmaps showing the footprints of RNA polymerase II and the directionality of transcription. The top row contains 227 genes that had the large CUT&RUN fragments directed towards downstream of the short-fragment peak (the first three column, positively skewed), and the bottom row contains 131 genes that had the large CUT&RUN fragments directed towards upstream of the short-fragment peak (the first three column, negatively skewed). Each gene is oriented from left to right in the heatmaps (i.e. genes encoded on the minus strand are all flipped horizontally). The heatmaps cover ±750 bp around the short-fragment peak of each gene. The fourth column onwards show the mNET-seq datasets from GSE60358. Each of the four datasets contains two heatmaps showing the nascent transcription from either the same strand (sense) or the opposite strand (antisense) of the gene.

We identified recurrent genes in which the large-size-class signals were either positively or negatively skewed (e.g. RBM6, SLC35B1, CCDC6 and HNRMPA0) (Fig. 3c and Online Resource 3). We anticipated that the skewed distribution of large-size-class reads correlates with the directionality of transcription from the promoter. To test this hypothesis, we turned to previously published mNET-seq data (Nojima et al. 2015) (Online Resource 4). The mNET-seq reads were aligned around the TSS of these genes (Fig. 3d). We observed that the genes were almost exclusively transcribed towards downstream (i.e. the same direction as the gene) from the promoter when the large-size-class Pol II footprints were positively skewed. On the other hand, bidirectional transcription was observed from the promoter of genes with negatively skewed Pol II footprints (Fig. 3d). These observations were confirmed by clustering genes by the skewness of the large-size-class signals and visualising the nascent transcription in these genes (Fig. 3e and Online Resource 5). In genes with positively skewed signals, sense transcription from the promoter (i.e. the same direction as the gene) was dominant. On the other hand, in genes with negatively skewed signals, abundant antisense transcription from the promoter was observed (Fig. 3e and Online Resource 5). Therefore, we conclude that the negatively skewed positioning of the large-size-class Pol II footprints is associated with the bidirectional transcription from the gene promoter (Nojima et al. 2015; Preker et al. 2008).

### Impact of formaldehyde fixation on “CUT&RUN on plate”

Formaldehyde fixation is often used to freeze transient protein interactions in biochemical assays. We tested if formaldehyde fixation can be implemented in “CUT&RUN on plate”. Cells were fixed with 1.5% formaldehyde before performing CUT&RUN. The resulting Pol II signals were mapped on the HOXA gene cluster region alongside the signals from unfixed cells (Fig. 4a). The formaldehyde fixation did not significantly affect the background cutting (Fig. 4b). However, the Pol II signal at the TSS (i.e. the maximum height of the Pol II peak) was ubiquitously reduced to some 10% to 30% of that in the “Unfixed” assays (Fig. 4c). We wish to see whether the observed signal reduction was due to the formaldehyde fixation or within the range of experimental variation. To this end, we compared CUT&RUN signals from eight independent assays (four “Unfixed” and four “Fixed” assays) (Fig. 4d). Difference in the signal intensity between unfixed and fixed samples were always greater than that between any two assays under the same condition (experimental variation) (Fig. 4e, cf. red and black symbols). Mean signal ratio between “Fixed” and “Unfixed” assays (log_10_0.7, n = 8) (Fig. 4e) indicates that the signal was reduced to about 20% on average in formaldehyde-fixed cells. It is possible that the formaldehyde fixation reduces the efficiency of chromatin cutting by pAG-MNase and/or affects the release of cleaved DNA fragments.

**Fig. 4.**
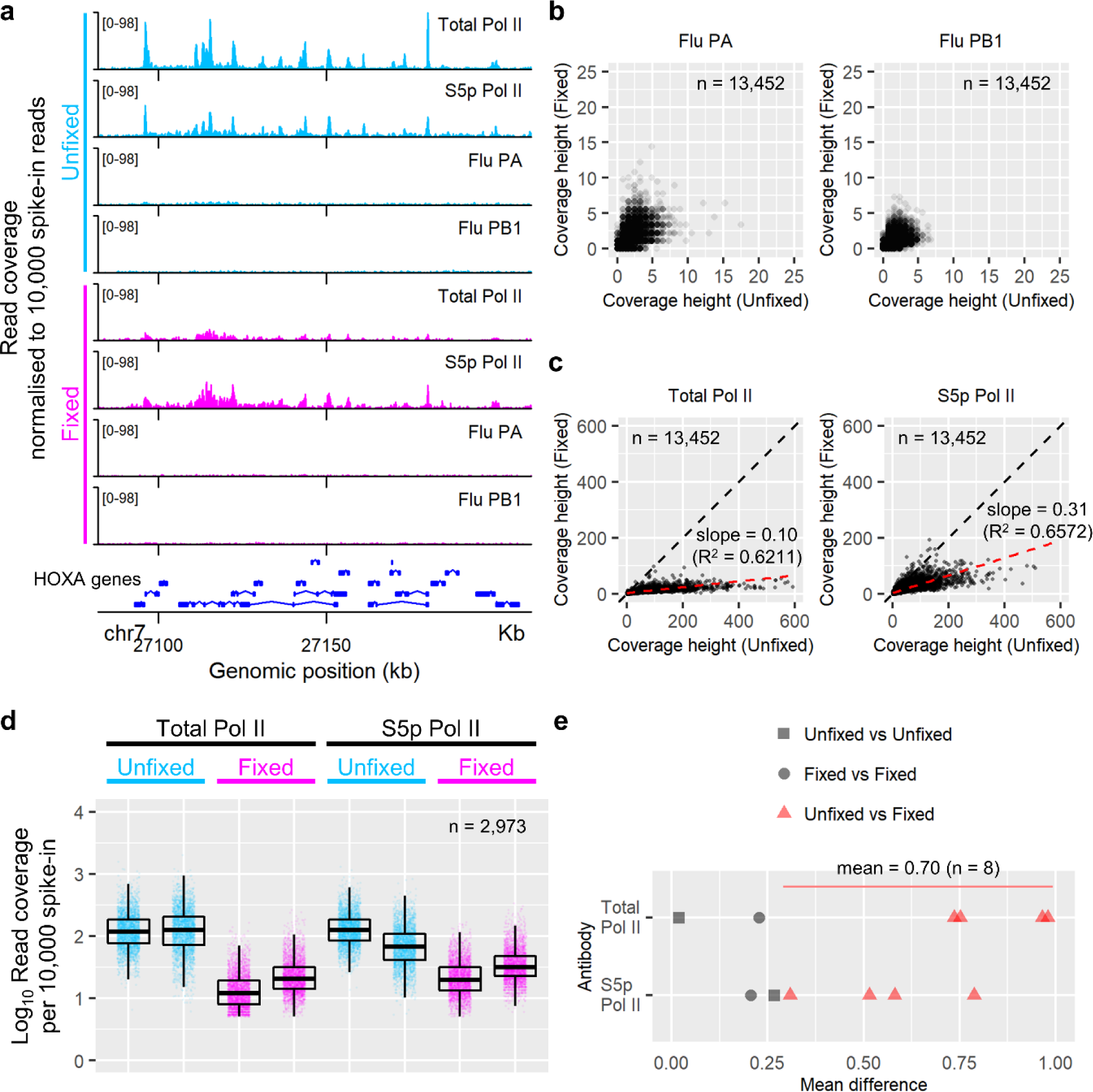
Impact of formaldehyde cell fixation on “CUT&RUN on plate”. (a) CUT&RUN signals of total fragments over the HOXA gene cluster (chr7 p15.2). Total Pol II, Millipore 05-623; and S5p Pol II, Abcam ab5131. Flu PA and Flu PB1 are negative controls obtained with irrelevant antibodies (influenza A PA and influenza A PB1, respectively). The vertical axis indicates the read coverage normalised to 10,000 spike-in reads. The scale is shown in brackets. Two colour codes are used to indicate whether cells were fixed with formaldehyde in the assays (light blue, “Unfixed”; and magenta, “Fixed”). (b) Comparison of the background cutting between the “Fixed” and “Unfixed” assays (13,452 genes). The maximum height of spike-in-normalised coverage within 1.5 kb around the TSS of each gene is shown. (c) Comparison of the Pol II signal intensity between the “Fixed” and “Unfixed” assays (13,452 genes). The maximum height of the spike-in-normalised Pol II peak (total reads) within 1.5 kb around the TSS is shown for each gene. The black dashed line indicates the diagonal line. The red dashed line shows a linear regression of these data points. The straight line was fitted to the data points and the parameters were estimated. (d) Spike-in-normalised Pol II signal intensity (maximum height of the total read coverage) around the TSS. The results of eight assays are presented (four for “Unfixed” and the other four “Fixed”). Genes with the signal above the background (peak height > 5) are shown (n = 2,973). The boxplot indicates 25%, 50% and 75% quantiles. (e) Mean difference of the Pol II intensity between two assays shown in panel (d). Data points comparing “Fixed” and “Unfixed” assays are highlighted in red; the mean of these eight datapoints (red triangles) is indicated.

### Summary

In this study, we have devised “CUT&RUN on plate” (CROP) for profiling protein-DNA interactions in adherent cells maintained in a multi-well cell culture plate. The current CUT&RUN protocol (Hainer and Fazzio 2019; Meers et al. 2019a; Skene et al. 2018) relies on Concanavalin A-coated magnetic beads to immobilise the cells before the targeted pAG-MNase cleavage. This immobilisation step introduces a range of shortcomings associated with the use of Concanavalin A-beads. For example, trypsinisation while harvesting the cells may alter the behaviour of DNA-binding proteins and lower the binding affinity of cells to Concanavalin A. Concanavalin A also stimulates certain types of cells. When the plasma membrane is removed to investigate the nuclei, Concanavalin A does not bind efficiently to the nuclear membrane. Finally, Concanavalin A-beads tend to aggregate when cells are attached. Our CROP protocol described here overcomes these key problems associated with the use of Concanavalin-A beads and may find widespread application. A fundamental limitation of the CROP method is that it is only applicable to adherent cells that are stably attached to the polystyrene surface of the culture plate throughout the protocol. In addition, it is crucial to seed the optimal number of cells, as well as to inspect the confluency of cells before an assay, to ensure that the cell count is consistent between assays for spike-in normalisation.

Using CUT&RUN, we profiled RNA polymerase II near the transcription start sites. Our observations suggest two distinct binding configurations of RNA polymerase II near the TSS. These configurations are: (1) RNA polymerase II at the TSS in the pre-initiation state where nucleosomes are depleted, and (2) paused RNA polymerase II upstream or downstream of the TSS where Pol II pausing is regulated by a higher-molecular complex (e.g. NELF) after transcription initiation (Fig. 5). The short-fragment (<120 bp) peak at the TSS were also observed for Serine 5-phosphyrylated Pol II (Fig. 2b), raising the possibility that some Pol II molecules detected by these short fragments are already released from the poised site, yet are unable to extend the transcription, which is known as abortive initiation (Goldman et al. 2009) The paused Pol II peaks were visible exclusively in the longer (>120 bp) fragments adjacent to the TSS. We reason that this is because Pol II is associated with a higher-molecular complex at the paused site (Vos et al. 2018), which protects the chromatin from the pAG-MNase cleavage. As a result, pAG-MNase is unable to produce shorter (<120 bp) fragments associated with the paused Pol II. We showed that the positioning of these large-size-class footprints was associated with the directionality of transcription from the gene promoter, demonstrating the biological significance of these CUT&RUN footprints.

**Fig. 5.**
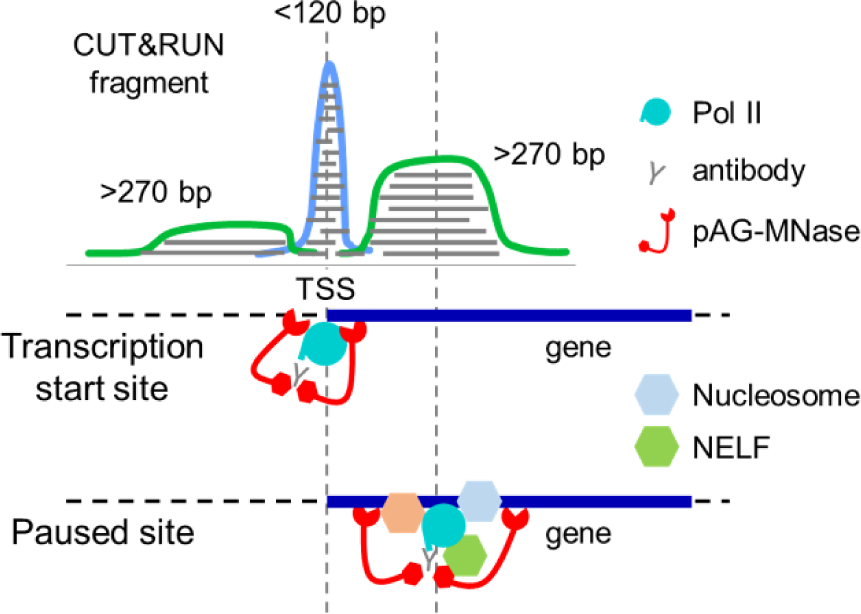
Working hypothesis on the Pol II footprints and Pol II pausing. In this working hypothesis, the short-size-class Pol II footprint indicates the transient positioning of poised Pol II, whereas the large-size-class Pol II footprint maps the promoter proximal pausing downstream of the transcription start site.

## Materials and Methods

### Cells

Human lung carcinoma cell line A549 was maintained in Dulbecco’s Modified Eagle Medium (Gibco, 11965-092) supplemented with 10% FBS (Gibco, 16140-071), 100 U/ml penicillin and 100 µg/ml streptomycin (Gibco, 15140-122).

### CUT&RUN in a cell culture plate

CUT&RUN (Skene and Henikoff 2017) was performed on a standard 24-well cell culture plate (TPP, 92012); cells were left attached to the bottom of the well throughout the protocol.

A549 was seeded in wells and grown to ∼70 to 90% confluency (∼0.5 million cells) at the time of an assay. Cells were washed with PBS and permeabilised with Perm buffer (0.1% Triton X-100 (Sigma, X100), 20 mM HEPES-KOH pH 7.5, 150 mM NaCl, 0.5 mM Spermidine (Sigma, S2626) and proteinase inhibitor (Roche, 04 693 132 001)) for 15 min at room temperature. After removing Perm buffer, cells were washed again with Perm buffer and incubated with an antibody (∼0.7 µg) in 150 µl Perm buffer per well for one hour at room temperature. The antibodies used in this study were: mouse anti-RNA polymerase II (Millipore, 05-623; clone CTD4H8; 0.75 µg per well); rabbit anti-RNA polymerase II (phospho S5) (Abcam, ab5131; polyclonal; 0.675 µg per well); rabbit anti-influenza A PA (Invitrogen, PA-32223; polyclonal; 1.1 µg per well); and rabbit anti-influenza A PB1 (Invitrogen, PA-34914; polyclonal; 0.86 µg per well). After incubation, unbound antibody was removed, and cells were washed twice with Perm buffer. Cells were then incubated with the recombinant pAG-MNase (Cell Signaling, 40366; 1:33 volume) in 150 µl Perm buffer per well for an hour at room temperature. Unbound pAG-MNase was removed, and cells were washed twice with Perm buffer. During the second wash of this stage, the plate was placed on ice-cold water. For pAG-MNase activation, 150 µl ice-cold Perm buffer containing 5 mM CaCl_2_ was dispensed per well, and the plate was incubated on ice-cold water for 30 min. The pAG-MNase digestion was halted by the addition of 50 µl 4× STOP solution (680 mM NaCl, 40 mM EDTA (Sigma, E5134), 8mM EGTA (AG Scientific, E-2491), 100 µg/ml Rnase A (Invitrogen, 12091021) and 0.1% Triton X-100). The plate was incubated at 37°C for 30 min to facilitate the release of digested DNA fragments from the nucleus. The supernatant was collected, and DNA was extracted using Dneasy Blood & Tissue Kit (Qiagen, 69504).

In addition to the native “CUT&RUN on plate” described above, two protocols for cell fixation with 1.5% formaldehyde were tested: (1) cells were fixed with 1.5% formaldehyde for 10 min prior to the cell permeabilisation with Perm buffer; and (2) cells were fixed following the permeabilisation (5 min in Perm buffer). Both protocols produced similar results. The supernatant collected at the end of the CUT&RUN protocol was supplemented with 1% SDS (final concentration) and incubated at 65°C overnight for de-crosslinking. The DNA was purified with QIAquick PCR Purification Kit (Qiagen, 28104).

### Pair-end sequencing and read alignment

DNA libraries were constructed using NEBNext Ultra II DNA Library Prep Kit for Illumina (NEB, E7645) with index primers (NEB, E7335, E7710 and E7600). Fragmented genomic DNA from *S. cerevisiae* (Cell Signaling, 40366) was used (10 pg per sample) as spike-in.

DNA libraries were sequenced (150 bp × 2) with NovaSeq 6000 (Illumina), and the initial 50 bases of each read were aligned to a concatenation of human reference genome (GRCh38) and *S. cerevisiae* (sacCer3) with a BWA algorithm (version 0.7.17) `bwa mem` (Li and Durbin 2009). Duplications of reads were marked with a Picard tool (version 2.18.9) `MarkDuplicates` (http://broadinstitute.github.io/picard). Only a minority of the aligned reads constituted the yeast genome, thereby confirming the successful recovery of DNA fragments cleaved by pAG-MNase. A subset of aligned reads was selected for the downstream analyses using Samtools (version 1.9) with the following commands: SAM flag 3 (read paired; read mapped in proper pair) (`samtools view -f 3`); mapping quality greater than 20 (`samtools view -q 20`); without tags “XA:Z:” and “SA:Z:” (`grep -v -e ‘XA:Z:’ -e ‘SA:Z:’`); and aligned in either one of the chromosomes chr1 to chr22, chrX or chrY (`samtools view chr1 chr2 … chrY`).

### Data analyses and visualisation

Aligned reads were fractionated by size (the 9th field of SAM) and stored in the BAM format. Read coverage was computed and stored in the bigWig format with deepTools’s (version 3.1.3) `bamCoverage` (Ramirez et al. 2016), normalised per 10,000 spike-in reads. The spike-in-normalised bigWig files were converted into the bedGraph format using `bigWigToBedGraph` (UCSC) and the read coverage was visualised with the R package Sushi (version 1.24.0) (Phanstiel et al. 2014) in R (version 3.6.2). A matrix which stores the read coverage over a given region of genes was computed using deepTools’s (version 3.3.2) `computeMatrix` with an annotation GCA_000001405.15_GRCh38_full_analysis_set.refseq_annotation (NCBI). A subset of genes that were separated from other genes by >2 kb was considered for this analysis; such genes were extracted using bedtools’s (version 2.26.0) (Quinlan and Hall 2010) `merge` and `intersect`. The matrix was visualised as a heatmap and a metaplot using the R package EnrichedHeatmap (version 1.16.0) (Gu et al. 2018).

### External mNET-seq data

Publicly available mNET-seq datasets (GSM1474225, GSM1474226, GSM1474228 and GSM1480181) were obtained from Gene Expression Omnibus (GSE60358) (Nojima et al. 2015). The original bigWig files were converted to the bedGraph format using bigWigToBedGraph (UCSC), and then the genome coordinates were converted to hg38 using liftOver (UCSC).

### Signal quantification and statistical tests

Pol II signal intensity were quantified by the maximum height of the spike-in-normalised read coverage within 1.5 kb around the annotated transcription start site (Fig. 3b and Figs. 4b-d). Local polynomial fitting to the short-fragment peak was performed using the R function `loess` (`span = 0.05`) (Fig. 3b). The signal in the upstream or downstream of the short-fragment peak (Fig. 3a) was quantified by the summation of signals in the 500 bp window stored in the matrix file from deepTools’s `computeMatrix` (Figs. 3a and c). Linear regression was performed using the R function `lm` to fit a straight line to the data points and estimate the parameters (Fig. 4c). Mean difference of two populations was calculated with the R function `wilcox.test` (Figs. 4d and e).

### Data reproducibility

The assays performed and analysed in this work are listed in Table 1. The source data for figure production are listed in Table 2.

**Table 1.** Assays performed in this study and data availability.

| Assay ID | Antibody <sup>a</sup> | Fixation |
| --- | --- | --- |
| 5-200331 | Total Pol II | Unfixed |
| 1-200518 | Total Pol II | Unfixed |
| 2-200518 | Total Pol II | Fixed <sup>b</sup> |
| 3-200518 | Total Pol II | Fixed <sup>c</sup> |
| 7-200331 | S5p Pol II | Unfixed |
| 7-200518 | S5p Pol II | Unfixed |
| 8-200518 | S5p Pol II | Fixed <sup>b</sup> |
| 9-200518 | S5p Pol II | Fixed <sup>c</sup> |
| 1-200820 | Total Pol II | Unfixed |
| 3-200820 | S5p Pol II | Unfixed |
| 4-200820 | Flu PA | Unfixed |
| 5-200820 | Flu PB1 | Unfixed |
| 1-1-200710 | Total Pol II | Fixed <sup>c</sup> |
| 1-3-200710 | S5p Pol II | Fixed <sup>c</sup> |
| 1-4-200710 | Flu PA | Fixed <sup>c</sup> |
| 1-5-200710 | Flu PB1 | Fixed <sup>c</sup> |

<sup>a</sup> Total Pol II, Millipore 05-623; S5p Pol II, Abcam ab5131
<sup>b</sup> Fixed after cell permeabilisation
<sup>c</sup> Fixed before cell permeabilisation

**Table 2.** Source data for figure production.

| **Figure** | **Source (Assay ID)** |
| --- | --- |
| Figs. 1b and c | Total Pol II: 1-200820<br>S5p Pol II: 3-200820<br>Flu PA (negative control): 4-200820<br>Flu PB1 (negative control): 5-200820 |
| Fig. 2a | Total Pol II: 5-200331<br>S5p Pol II: 7-200518 |
| Online Resource 1<br>(Supplementary to Fig. 2a) | Total Pol II: 1-200518<br>S5p Pol II: 7-200331 |
| Fig. 2b | Total Pol II: 5-200331<br>S5p Pol II: 7-200518 |
| Online Resource 2<br>(Supplementary to Fig. 2b) | Total Pol II: 1-200518<br>S5p Pol II: 7-200331 |
| Fig. 2c | Total Pol II: 5-200331 (Assay 1) and 1-200518 (Assay 2)<br>S5p Pol II: 7-200331 (Assay 1) and 7-200518 (Assay 2) |
| Figs. 3b and c | 5-200331 |
| Online Resource 3<br>(Supplementary to Fig. 3c) | Total Pol II: 1-200518<br>S5p Pol II (left): 7-200331 (left) and 7-200518 (right) |
| Fig. 3d | Total Pol II: 5-200331 (top) and 1-200518 (bottom)<br>S5p Pol II: 7-200331 (top) and 7-200518 (bottom) |
| Fig. 3e | 5-200331 |
| Online Resource 5<br>(Supplementary to Fig. 3e) | Total Pol II (1-200518)<br>S5p Pol II: 7-200331 (top) and 7-200518 (bottom) |
| Figs. 4a-c | Total Pol II: 1-200820 (Unfixed) and 1-1-200710 (Fixed)<br>S5p Pol II: 3-200820 (Unfixed) and 1-3-200710 (Fixed)<br>Flu PA: 4-200820 (Unfixed) and 1-4-200710 (Fixed)<br>Flu PB1: 5-200820 (Unfixed) and 1-5-200710 (Fixed) |
| Figs. 4d and e | Total Pol II (Unfixed): 5-200331 and 1-200518<br>Total Pol II (Fixed): 3-200518 and 2-200518<br>S5p Pol II (Unfixed) :7-200331 and 7-200518<br>S5p Pol II (Fixed): 9-200518 and 8-200518 |

## Abbreviations

CUT&RUN: Cleavage Under Targets and Release Using Nuclease
pAG-MNase: Protein A/G-fused micrococcal nuclease
ChIP: chromatin immunoprecipitation
Pol II: RNA polymerase II
TSS: transcription start site
S5p Pol II: Serine 5-phosphorylated RNA polymerase II
mNET-seq: mammalian native elongating transcript-sequencing

## Acknowledgements

We thank Liew Jun Mun, University of Cambridge, for reviewing the manuscript to enhance the readability. We also thank Prof. Charles R. M. Bangham, Imperial College London, for reviewing the manuscript and critical comments. Illumina NovaSeq 6000 sequencing was performed at Centre for PanorOmic Sciences (CPOS) Genomics Core, LKS Faculty of Medicine, The University of Hong Kong. This study was supported in part by the Research Grants Council of the Hong Kong SAR (17119618, 17107019 and C7027-16G).

## Declarations

### Ethics approval

Not applicable

### Consent for publication

Not applicable

### Authors’ contributions

MM: Conceptualization, Methodology, Validation, Formal analysis, Investigation, Data Curation, Visualization, Writing - original draft preparation, Writing - review and editing HC: Resources, Project administration, Funding acquisition, Writing - review and editing

## Supplementary information

**Online Resource 1.**
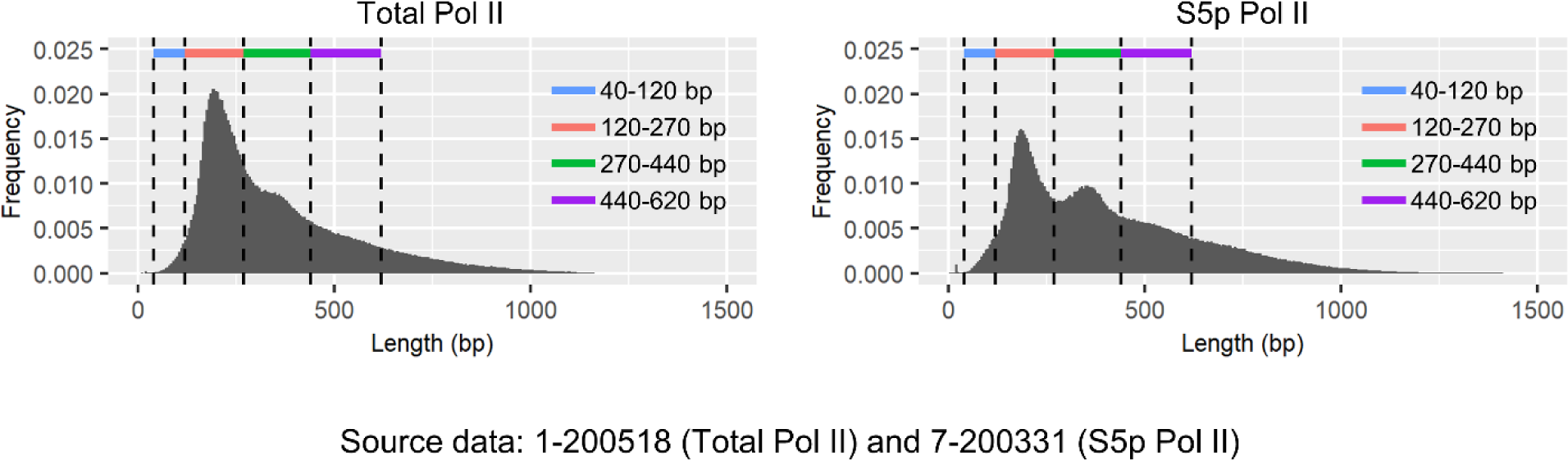
Length distribution of aligned reads (Supplementary to Fig. 2a)

**Online Resource 2.**
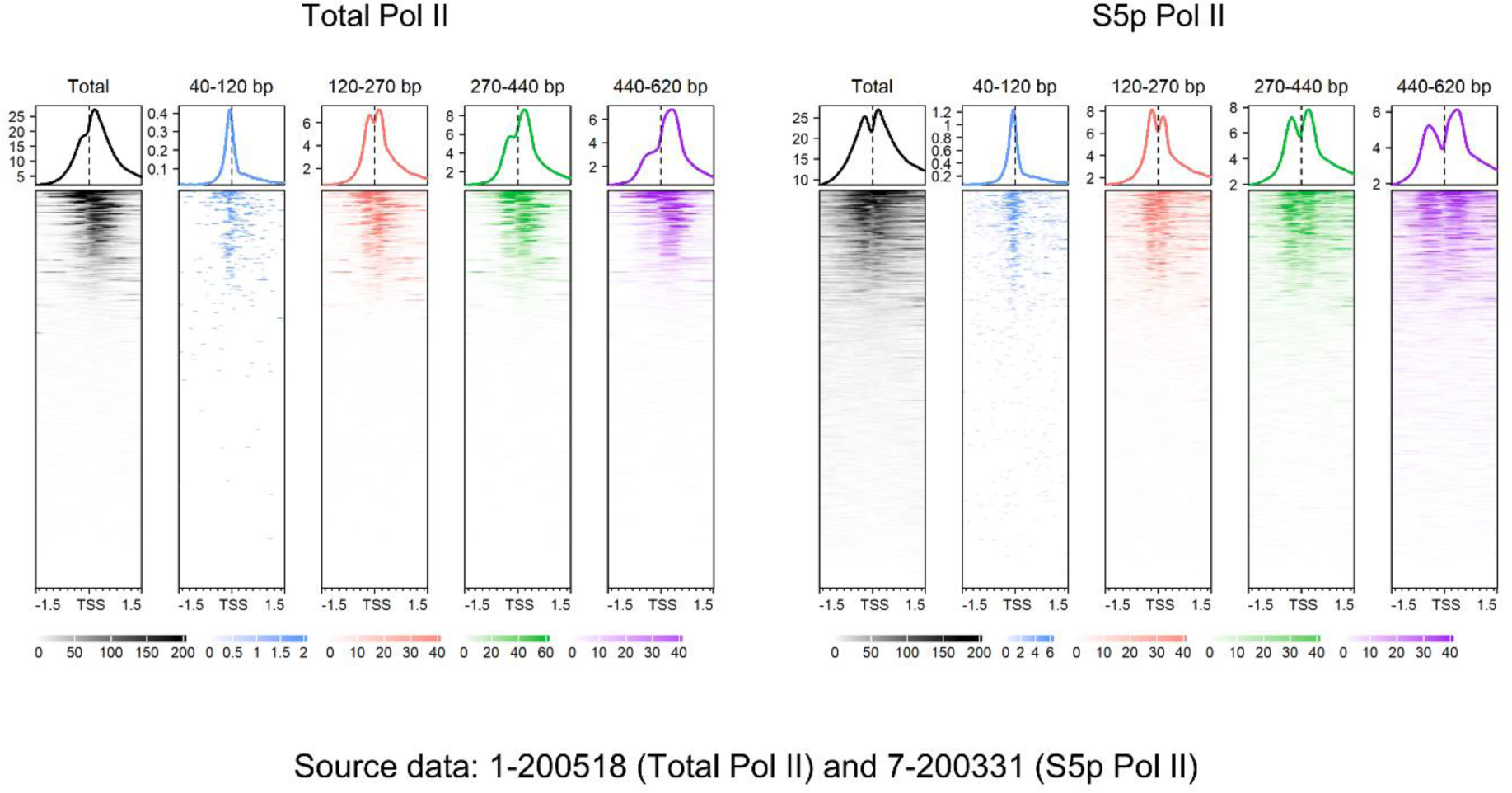
CUT&RUN fragments and Pol II footprints (Supplementary to Fig. 2b)

**Online Resource 3.**
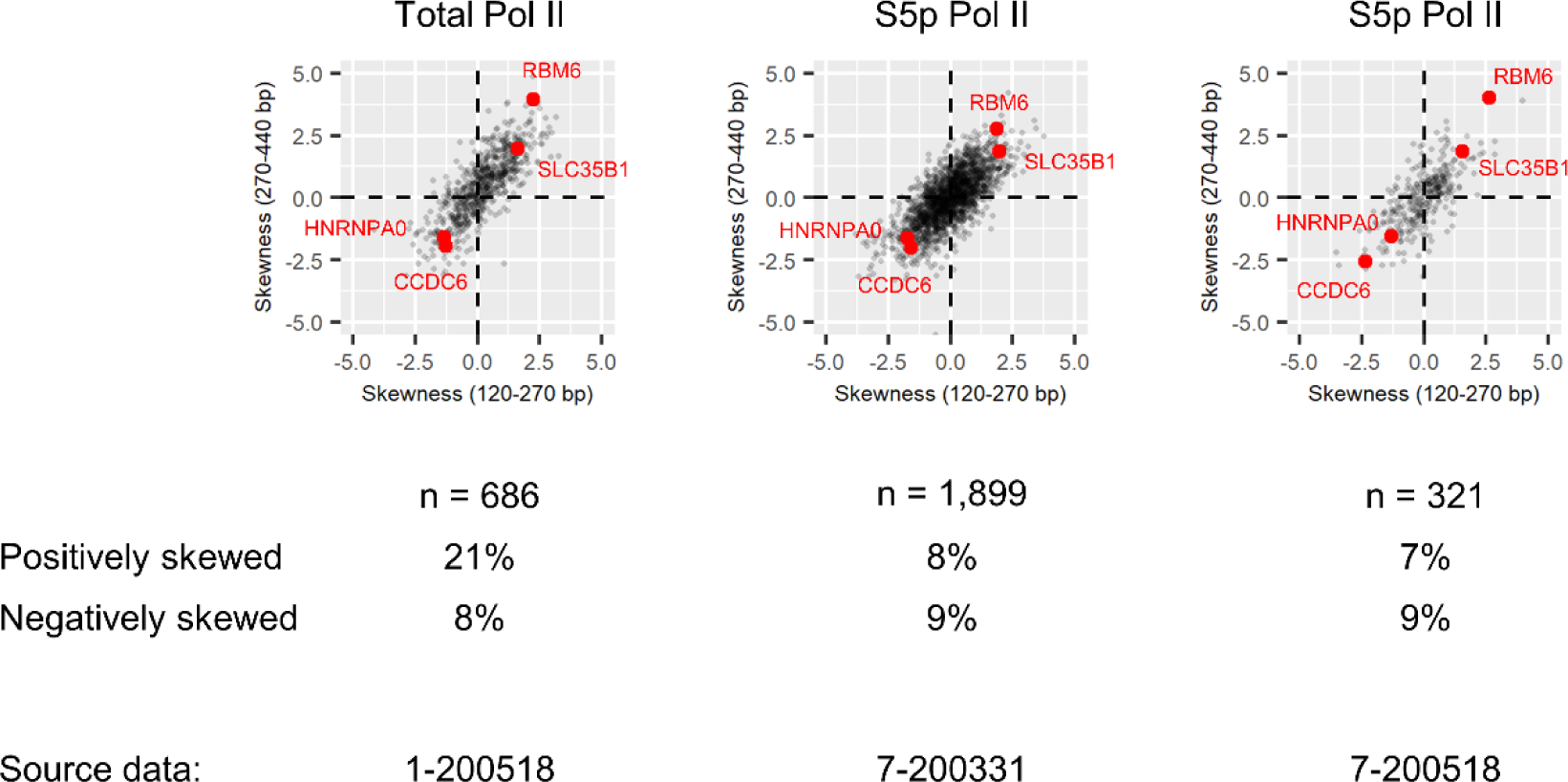
Skewness of the large-size-class Pol II footprints (Supplementary to Fig. 3c)

**Online Resource 4.**
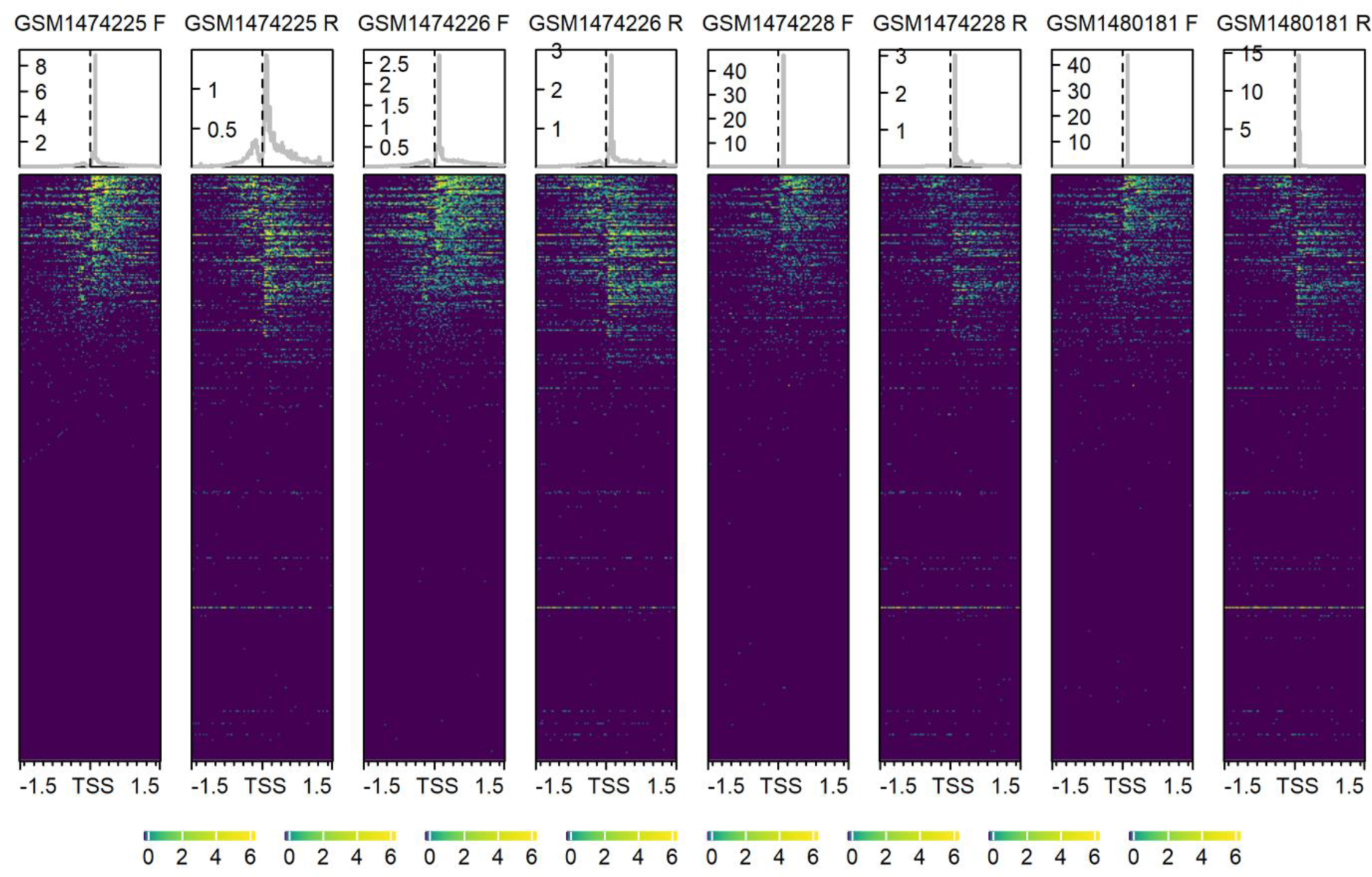
mNET-seq datasets used in this study Publicly available mNET-seq data (GSE60358). The heatmap shows the nascent transcription on the plus (F) or the minus (R) strand within ±1.5 kb around the annotated transcription start site of each gene. The original data was lifted to hg38. The axis at the bottom indicates the genomic position (kb) around the transcription start site. RefSeq genes that are separated from other genes by >2 kb are presented (n = 13,452).

**Online Resource 5.**
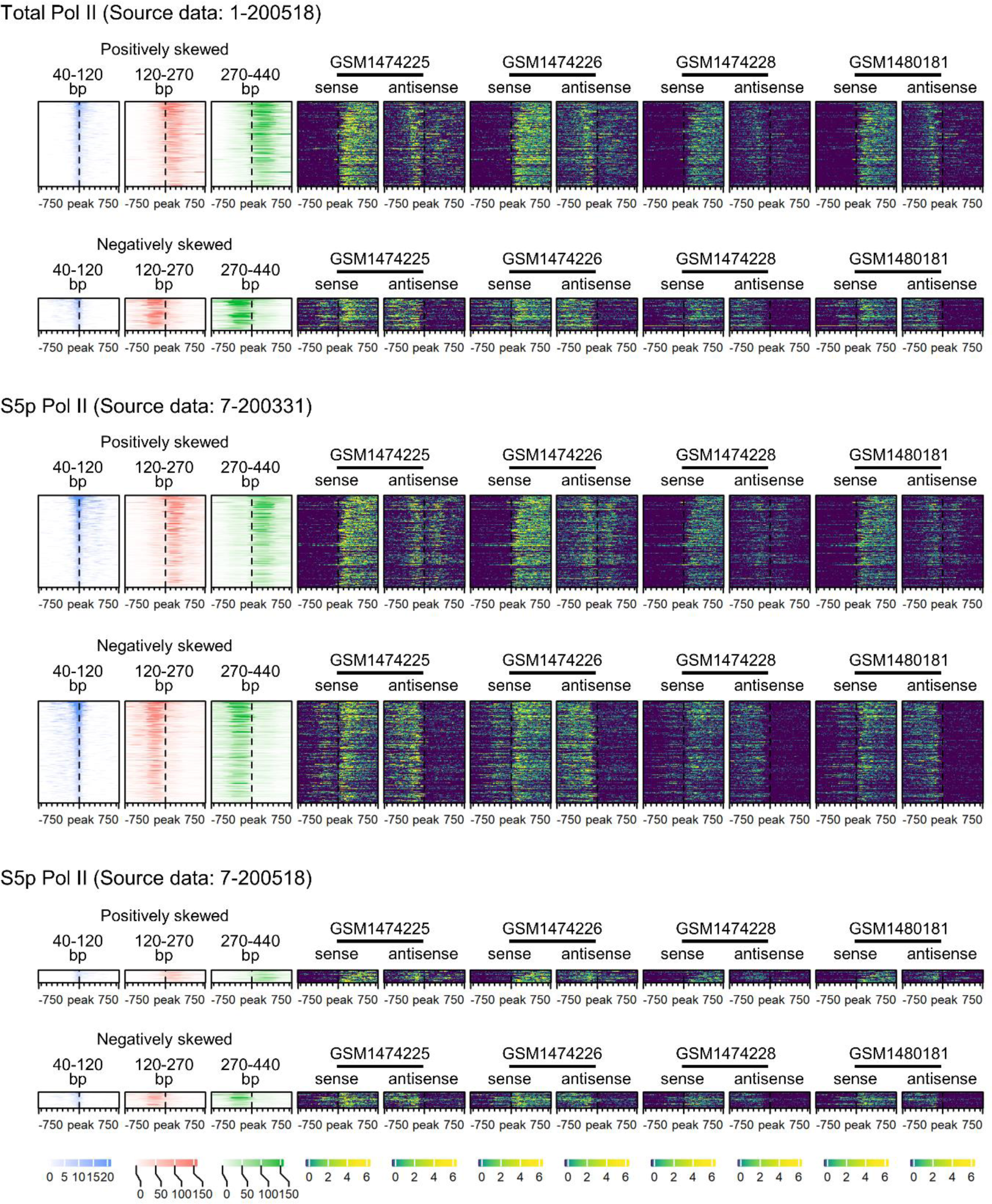
Skewed Pol II footprints and bidirectional transcription (Supplementary to Fig-3e)

